# EpiCompare: R package for the comparison and quality control of epigenomic peak files

**DOI:** 10.1101/2022.07.22.501149

**Authors:** Sera Choi, Brian M. Schilder, Leyla Abbasova, Alan E. Murphy, Nathan G. Skene

## Abstract

**Summary:** EpiCompare combines a variety of downstream analysis tools to compare, quality control and benchmark different epigenomic datasets. The package requires minimal input from users, can be run with just one line of code and provides all results of the analysis in a single interactive HTML report. EpiCompare thus enables downstream analysis of multiple epigenomic datasets in a simple, effective and user-friendly manner.

**Availability and Implementation:** EpiCompare is available on Bioconductor (≥ v3.15):

https://bioconductor.org/packages/release/bioc/html/EpiCompare.html

All source code is publically available via GitHub:

https://github.com/neurogenomics/EpiCompare

Documentation website

https://neurogenomics.github.io/EpiCompare

EpiCompare DockerHub repository:

https://hub.docker.com/repository/docker/neurogenomicslab/epicompare

## Introduction

Epigenetic processes are crucial regulators of gene expression and transcriptional activity (Allis & Jenuwein, 2016). There is an increasing interest towards understanding disease mechanisms with epigenetic factors, especially in cancer (Cheng et al., 2019), autoimmune diseases (Mazzone et al., 2019) and brain disorders (Hannon et al., 2019; Roussos et al., 2014). In response to this, a variety of novel epigenomic profiling technologies have emerged in recent years (Cazaly et al., 2019; Mehrmohamadi et al., 2021). Yet, how the performance of these new methods compare with traditional approaches and how they differ from one another is largely unknown. Therefore, we need a way of systematically comparing and analysing epigenomic data generated by different methods to contrast their strengths and weaknesses. This would help researchers to choose the appropriate method when designing epigenomic studies and to establish the optimal experimental protocols and data analysis workflows.

Typically, epigenomic data analysis consists of two parts: (1) data processing, where sequences are mapped and peaks are called; and (2) downstream analysis, where peaks are visualised and annotated. There have been movements to standardise and simplify data preprocessing steps through workflow-based pipelines (Ewels et al., 2020) such as *nf-core/chipseq* (Patel et al., 2021) and *nf-core/cutandrun (Cheshire et al., 2022)*, which require just one line of code to run. However for the downstream analysis, the tools are currently scattered in many different packages and platforms, with some requiring idiosyncratic input formats. This makes the latter part of the analysis challenging and time-consuming, especially for those with little or no computational experience.

To address these issues, we introduce EpiCompare, a Bioconductor (Huber et al., 2015) R package for the comparison and quality control of epigenomic data. EpiCompare is able to perform a variety of downstream analysis on multiple epigenomic datasets simultaneously, which can be executed with just one line of code. Some of the main functionalities include precision-recall and functional annotations, which help to assess the extent of overlapping peaks between files and to check if peaks annotate to the same genomic features. The package also generates a single report collating all results of the analysis into a single interactive report file, making it easy for users to view and interpret the results.

## Implementation

EpiCompare was implemented using the R programming language (v4.2) (R Core Team, 2021) in accordance with all Bioconductor (Huber et al., 2015) coding and documentation standards. In addition, every time an update is pushed to EpiCompare, extensive checks and unit tests are automatically launched on three OS platforms (Unix, Mac, Window) via continuous integration workflows using both GitHub Actions and Bioconductor. The package EpiCompare can launch all analyses using a single master function (eponymously named *EpiCompare*). Users need only to supply peak files of interest as a named list of GenomicRanges objects (Lawrence et al., 2013) or as paths to BED files to be automatically imported as GenomicRanges. It also includes several Boolean parameters, allowing users to decide which analyses to perform. When the function is executed, it parses the parameters into an R markdown file, which is ultimately rendered into an HTML document.

First, *EpiCompare* runs three quality control and standardisation checks on all input peak files. The first step involves removal of blacklisted genomic regions containing well-known irregular or anomalous signals (Amemiya, Kundaje & Boyle, 2019). Filtering out these peaks is recommended for quality measures and thus, *EpiCompare* requires that users specify a ‘blacklist’ peak file. The second control step uses BRGenomics (v1.1.3) *tidyChromosomes* (DeBerardine, 2022) feature to remove peaks that are found in non-standard or mitochondrial chromosomes. Lastly, the final check ensures that all input peak files are based on the same reference genome build. Users must specify the genome build used to generate the peak files and if needed, *EpiCompare* uses rtracklayer (v1.56.0) *liftOver* (Lawrence, Gentleman & Carey, 2009) function to translate the genomic coordinate of peak files across builds.

## Usage

EpiCompare can be installed on any Unix, Mac, or Windows OS using BiocManager. Alternatively, EpiCompare can be installed using its dedicated Docker or Singularity container (hosted on DockerHub), which greatly alleviates common challenges with installation and reproducibility. Once EpiCompare is installed, the package can be used with a single line of code or one function call (*EpiCompare*). The function requires two inputs: a list of peak files of interest and a ‘blacklist’ peak file containing genomic regions of irregular signals. All peak data can be specified as GenomicRanges objects or as paths to BED files. To ensure that genome builds of peak files agree, users must also state the genome build that was used to generate the peak, reference and blacklist files, which can be supplied as a single genome build (e.g. *genome_build=“hg19”*) or a named list of mixed genome builds (e.g. *genome_build=list(peakfiles=“hg19”, reference=“hg38”, blacklist=“hg38”)*). In addition to human genome builds (hg19, GRCh38), *EpiCompare* can ingest and/or output files aligned mouse (mm9, mm10) genome builds using interspecies chain files.

In addition, *EpiCompare* offers a suite of analysis tools and plot options to choose from, allowing users to tailor their downstream analysis of epigenomic data. The two optional inputs include reference peak file and duplicate summary outputs from Picard (Broad Institute, 2019). The plot options are summarised in **Table 1**. Once all analyses are complete, the rendered HTML file can be automatically launched in any web browser, or within Rstudio (RStudio Team, 2022). All data and plots produced by the analyses are also stored in a subfolder called “EpiCompare_files”.

**Table 1.** Summary of plot options in *EpiCompare*

| Plot | Description |
| --- | --- |
| Upset Plot | Upset plot of the number of overlapping peaks between files. |
| Stat Plot | Box plot showing the distribution of statistical significance (q-values) of sample peaks that are overlapping and non-overlapping with the reference peak file. |
| Precision-recall plot | Computes precision-recall curves across different peak strength thresholds. Metadata columns to be used for thresholding can be automatically inferred based on known relevant columns generated by SEACR, MACS2/3 and HOMER. |
| Correlation plot | Standardises all peak files by rebinning them into tiles of a user-defined width across the genome, and then compute pairwise correlation statistics between all peak files. |
| ChromHMM Plot | Heatmap of ChromHMM(6) annotation of peaks. |
| ChIPseeker Plot | Bar chart of ChIPseeker(7) annotation of peaks. |
| Enrichment Plot | Dot plot of KEGG pathway and GO enrichment analysis of peaks. |
| TSS Plot | Peak frequency around (+/-3000bp) transcriptional start site. |

EpiCompare can also call consensus peaks from groups of peak files via the function *compute_consensus_peaks*. Multiple methods for calling consensus peaks are offered, including a fast but simplistic overlap strategy (*method=“granges”*), and a slower but more accurate strategy that incorporates modelling of peak distributions (*method=“consensusseeker”*) (Samb et al., 2015). This can be helpful as a pre-step for reducing the number of samples being input to *EpiCompare*, and making files more comparable to “replicated peaks” files in databases like ENCODE.

All of the core functions used internally by the main function *EpiCompare* are exported so that they can be used in custom workflows as they may be more generally useful to the bioinformatics community (e.g. *compute_consensus_peaks, gather_files, plot_precision_recall, compute_corr, rebin_peaks, overlap_heatmap, plot_enrichment liftover_grlist*). See here for documentation on all exported functions: https://neurogenomics.github.io/EpiCompare/reference

## Output

*EpiCompare* generates an interactive HTML report containing all results of the analysis. The exact code used to generate each section is embedded within the report (collapsed by default). The report is organised into three parts: General Metrics, Peak Overlap and Functional Annotation. All sections are easily navigated by an interactive table of contents. The General Metrics section presents information on individual peak files, including the number of peaks, percentage of peaks in blacklisted regions and non-standard chromosomes, distribution of peak width, and duplication rate of mapped fragments. The Peak Overlap section provides the frequency, percentage, statistical significance, precision-recall and correlation of overlapping and non-overlapping peaks between the sample and reference peak files. Finally, the Functional Annotation section contains the functional annotation of peaks. These are ChromHMM (Ernst & Kellis, 2017), ChIPseeker (Yu, Wang & He, 2015), enrichment analysis (KEGG pathway and GO) and the frequency of peaks around the transcriptional start site.

To demonstrate the functionalities, we used EpiCompare to contrast the profiling of open chromatin regions of human K562 cells using ATAC-seq and DNase-seq. Several of the figures included in the report can be seen in Figure 1 (see https://neurogenomics.github.io/EpiCompare/inst/report/EpiCompare_example.html for the full report). The two ATAC-seq (ENCFF558BLC and ENCFF333TAT) and DNase-seq (ENCFF274YGF and ENCFF185XRG) datasets were obtained from ENCODE (ENCODE Project Consortium, 2012). Using EpiCompare, we can see a difference between the two methods, especially in ChromHMM annotations and precision-recall plot (Figure 1d&1e).

**Figure 1.**
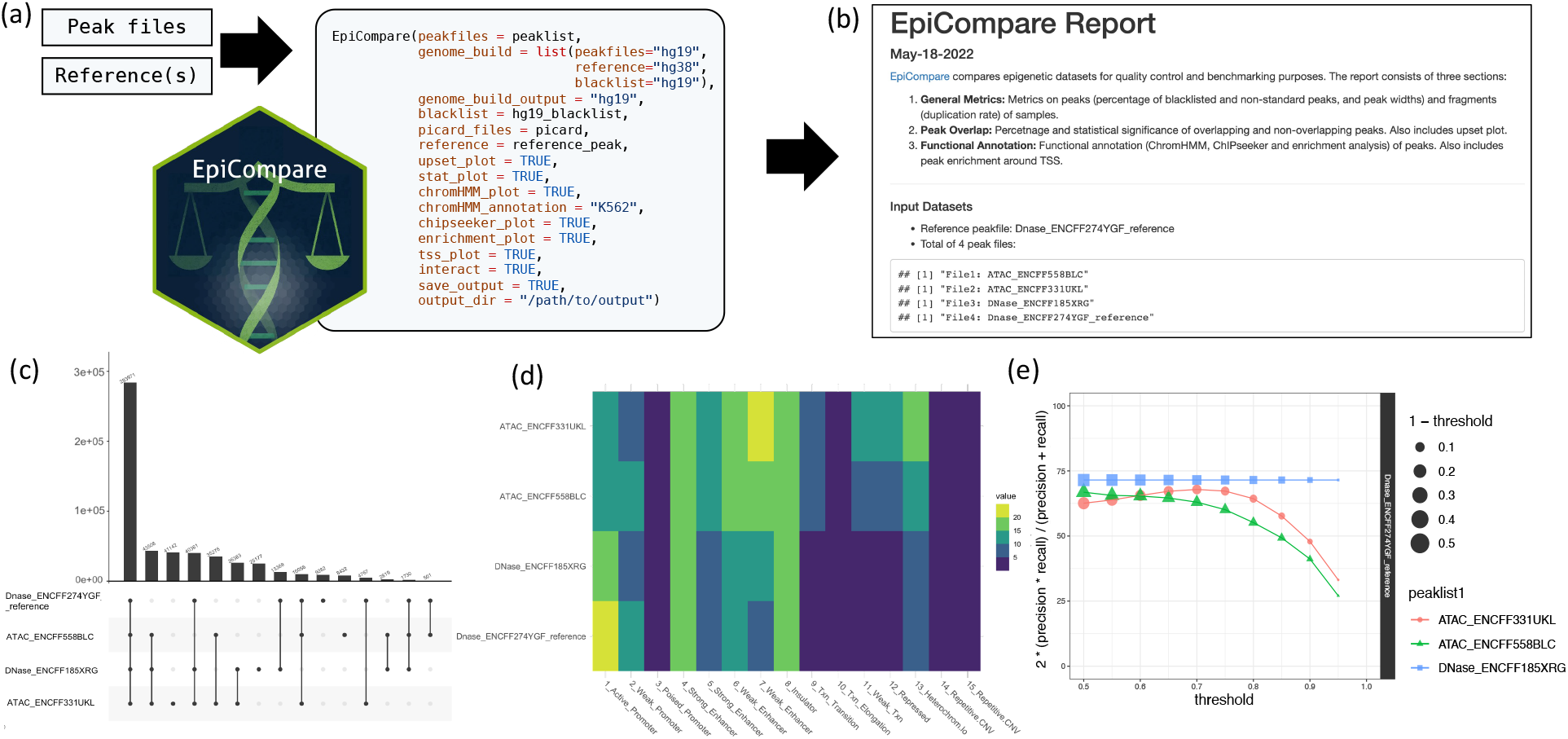
Flowchart demonstrating the use of *EpiCompare*. This example compares the open chromosome regions of human K562 cells profiled using ATAC-seq and DNase-seq. (a) Peak files are input into the master function (*EpiCompare*). (b) The function outputs an HTML report containing all results of the analysis. (c) Upset plot showing the number of overlapping peaks between peak files. (d) ChromHMM annotation of peak files. (e) Plot showing the precision-recall score across the peak calling stringency thresholds.

## Conclusion

Here, we presented EpiCompare, a Bioconductor R package for the comparison and quality control of epigenomic data. The package offers a selection of downstream analysis tools, enables processing of multiple epigenomic datasets in parallel and allows users to tailor their analyses. All of this can be executed with just one R function with minimal input from users, making the usage less demanding for those with little computational experience. Lastly, it generates a single report containing all results of the analysis, providing a simple, efficient and user-friendly way of comparing epigenomic datasets. EpiCompare will continue to be optimised and enhanced over time, with new features such as API access to thousands of peak files stored on public databases already underway.

## Acknowledgements

We like to thank Sarah Marzi, Alexi Nott and Di Hu for contributing to the discussions of early results and for suggesting new functionalities of EpiCompare throughout the development process.

